# VIP interneurons regulate cortical size tuning and visual perception

**DOI:** 10.1101/2023.03.14.532664

**Authors:** Katie A. Ferguson, Jenna Salameh, Christopher Alba, Hannah Selwyn, Clayton Barnes, Sweyta Lohani, Jessica A. Cardin

## Abstract

Local cortical circuit function is regulated by diverse populations of GABAergic interneurons with distinct properties and extensive interconnectivity. Inhibitory-to-inhibitory interactions between interneuron populations may play key roles in shaping circuit operation according to behavioral context. A specialized population of GABAergic interneurons that co-express vasoactive intestinal peptide (VIP-INs) are activated during arousal and locomotion and innervate other local interneurons and pyramidal neurons. Although modulation of VIP-IN activity by behavioral state has been extensively studied, their role in regulating information processing and selectivity is less well understood. Using a combination of cellular imaging, short and long-term manipulation, and perceptual behavior, we examined the impact of VIP-INs on their synaptic target populations in the primary visual cortex of awake behaving mice. We find that loss of VIP-IN activity alters the behavioral state-dependent modulation of somatostatin-expressing interneurons (SST-INs) but not pyramidal neurons (PNs). In contrast, reduced VIP-IN activity disrupts visual feature selectivity for stimulus size in both populations. Inhibitory-to inhibitory interactions thus directly shape the selectivity of GABAergic interneurons for sensory stimuli. Moreover, the impact of VIP-IN activity on perceptual behavior varies with visual context and is more acute for small than large visual cues. VIP-INs thus contribute to both state-dependent modulation of cortical circuit activity and sensory context-dependent perceptual performance.

## Introduction

The function of local circuits in the neocortex is shaped by the activity of a diverse set of GABAergic interneurons with distinct intrinsic properties, connectivity, and synaptic dynamics (Fishell and Rudy, 2011; Rudy et al., 2011; Pfeffer et al., 2013; Tremblay et al., 2016). Recent work has highlighted the role of a specialized population of vasoactive intestinal peptide-expressing interneurons (VIP-INs) in the state-dependent regulation of cortical activity (Fu et al., 2014; Pakan et al., 2016; Dipoppa et al., 2018). VIP-INs are depolarized by acetylcholine (Porter et al., 1999; Fu et al., 2014; Askew et al., 2019; Gasselin et al., 2021; Ren et al., 2022) and active during periods of high arousal, behaviorally relevant input (Pi et al., 2013; Szadai et al., 2022), and locomotion (Fu et al., 2014; Pakan et al., 2016; Batista-Brito et al., 2017; Dipoppa et al., 2018). Their influence on local circuit operations may thus be selectively exerted according to behavioral context.

The influence of VIP-INs on the surrounding circuit is largely thought to arise through their robust inhibition of another population of GABAergic interneurons that co-express the peptide somatostatin (SST-INs) (Pfeffer et al., 2013; Karnani et al., 2014; Karnani et al., 2016a). VIP-IN inhibition of SST-INs leads to disinhibition of local excitatory pyramidal neurons (PNs), causing amplification of PN activity (Lee et al., 2013; Pi et al., 2013; Fu et al., 2014; Zhang et al., 2014; Jackson et al., 2016; Karnani et al., 2016b; Kuchibhotla et al., 2017; Heintz et al., 2022). VIP-INs also receive inhibition from SST-INs and parvalbumin-expressing interneurons (PV-INs) (Hioki et al., 2013; Pfeffer et al., 2013; Jiang et al., 2015) and directly inhibit both local PV-INs (Walker et al., 2016) and the dendrites of local PNs (Acsady et al., 1996a; Acsady et al., 1996b; Tyan et al., 2014; Garcia-Junco-Clemente et al., 2017; Chiu et al., 2018). Inhibitory-to-inhibitory synaptic interactions among VIP-, SST-, and PV-INs may maintain a balance between inhibition and disinhibition, and likely play a role in stabilizing the operation of cortical circuits (Tsodyks et al., 1997; Ozeki et al., 2009; Litwin-Kumar et al., 2011; Litwin-Kumar et al., 2016; Millman et al., 2020; Sadeh and Clopath, 2021).

Although the state-dependent modulation of VIP-INs has been extensively characterized (Fu et al., 2014; Pakan et al., 2016; Munoz et al., 2017; Dipoppa et al., 2018) their impact on cortical sensory encoding is less well understood. The complex interactions among cortical interneurons may permit sensory context-dependent engagement of distinct modes of inhibitory modulation of local cortical circuits (Kuchibhotla et al., 2017). In mouse primary visual cortex, where interneuron response properties have been extensively studied, VIP-INs have small receptive fields (Mesik et al., 2015; Dipoppa et al., 2018). Recent work found that VIP-INs respond primarily to small, low-contrast visual stimuli and are suppressed in response to larger, more salient stimuli (de Vries et al., 2020; Millman et al., 2020). VIP-IN inhibition of SST-INs depends on visual context and is robust when stimulus center and surround differ but may be reduced when center and surround match (Keller et al., 2020). Moreover, manipulation of VIP-IN activity can modulate overall levels of cortical activity (Jackson et al., 2016) and some visual response properties of PNs (Ayzenshtat et al., 2016). However, the contributions of interactions between VIP and SST interneurons to visual tuning of local PNs remains unclear. Furthermore, the degree to which VIP-IN activity regulates visual perception is unknown. Here, we use short and long-term manipulation of VIP-INs to examine their role in shaping both the behavioral state-dependent visual feature selectivity of cortical SST-INs and PNs and visual perceptual behavior.

## Results

### Targeted removal of VIP interneurons in V1

VIP-INs are activated at the onset of locomotion and other high arousal states, suggesting that they contribute to state-dependent modulation of local circuit activity. We tested the role of VIP-INs in regulating the state modulation of local neurons, using mouse primary visual cortex (V1) as a model of local circuit function. VIP-INs synapse on local somatostatin-expressing interneurons (SST-INs) (Pfeffer et al., 2013; Neske and Connors, 2016; Walker et al., 2016), which provide dendritic inhibition to excitatory pyramidal cells (PNs) (Kapfer et al., 2007; Silberberg and Markram, 2007; Ma et al., 2010). However, VIP-INs also target other IN populations and directly inhibit PN dendrites (Acsady et al., 1996a; Acsady et al., 1996b; Tyan et al., 2014; Garcia-Junco Clemente et al., 2017; Chiu et al., 2018). Thus, to identify the impact of VIP inhibition on downstream targets in the local circuit, we used 2-photon imaging to measure the activity of SST-INs and PNs in control and VIP- ablated animals (see Methods). We selectively caused apoptotic cell death of VIP-INs via expression of a conditional viral construct carrying a genetically engineered caspase-3 (Yang et al., 2013) (Fig. 1A). Cell type specific expression of caspase caused rapid cell death in >75% of VIP-INs in V1 within two weeks of viral injection (SFig. 1A-D).

**Figure 1.**
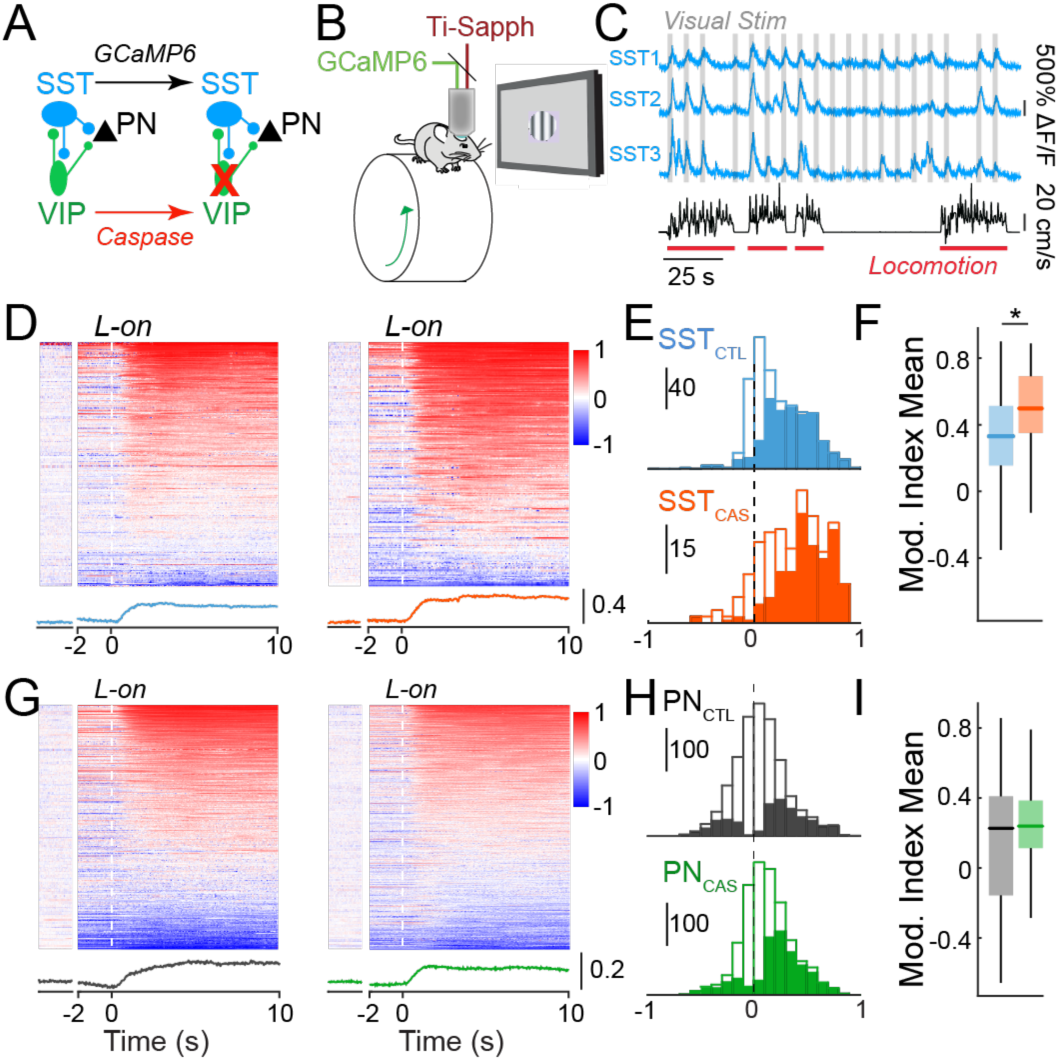
VIP interneuron ablation selectively disrupts state-dependent activity of SST interneurons. (A) Cre-dependent expression of Caspase-3 causes cell death in VIP-INs. GCaMP6 is expressed in SST-INs or PNs in each experiment. (B) Schematic of the in vivo 2-photon imaging configuration. (C) Ca2+ traces of three example SST-INs (blue) recorded during the presentation of visual stimuli (grey) and wheel speed tracking (black) to identify locomotion bouts (red). (D) Modulation of activity around locomotion onset (L-on), calculated as an index value, of SST-INs in control animals (left) and VIP ablation animals (right). Modulation during periods of sustained quiescence (Q) (see Methods) is shown to the left for comparison. Average modulation trace shown below heatmaps for controls (blue) and VIP ablation animals (orange). (E) Histogram of modulation indices of all SST-INs in control (SST_CTL_, blue; n = 603 cells in 7 mice) and VIP ablation animals (SST_CAS_, orange; n = 283 cells in 5 mice). Solid bars indicate cells showing significant modulation at p<0.05 (shuffle test). (F) Box plot of locomotion modulation indices in E. Central mark indicates the median, and whiskers indicate 25th and 75th percentiles. (G-I) Same analysis as in D-F but for PNs in control (PN_CTL_, black; n = 1694 cells in 6 mice) and VIP- IN ablation animals (PN_CAS_, green; n = 1623 cells in 6 mice). *p<0.05, linear mixed effects model.

We expressed the calcium indicator GCaMP6 in SST-INs and PNs in layer 2/3 of V1 and imaged each population in awake, head-fixed adult control and VIP ablation mice (Figs. 1, S1) across waking behavioral states (Fig. 1B-C). In controls, the majority of SST-INs exhibited increases in activity with high arousal, marked by the onset of locomotion (Fig.1D, E) (Fu et al., 2014; Pakan et al., 2016; Dipoppa et al., 2018), whereas PNs showed a more diverse profile (Fig. 1G,H) (Niell and Stryker, 2010; Vinck et al., 2015). In the absence of VIP-INs, state dependent modulation of spontaneous SST-IN activity was significantly enhanced (Fig. 1D-F), suggesting that VIP inhibition of SST-INs normally regulates the spontaneous activity of these cells. In contrast, loss of VIP-INs had no impact on the state-dependent regulation of PN activity (Fig. 1G-I). Together, these data suggest that loss of VIP-INs has a strong impact on the regulation of SST-INs, but not PNs, by behavioral state.

Experimental data and computational modeling of cortical circuits suggest that inhibition plays a role in stabilizing local networks (Ozeki et al., 2009; Litwin-Kumar et al., 2011; Litwin-Kumar et al., 2016; Sadeh and Clopath, 2021) and coordinating patterns of activity (Cardin et al., 2009; Cardin, 2018; Veit et al., 2023). To examine the impact of VIP inhibition on the coordinated activity of local SST-INs and PNs, we presented repeated high-contrast visual stimuli and measured noise correlations (Cohen and Kohn, 2011) within each population. Loss of VIP-INs led to increased modulation of noise correlations among SST-INs, but not PNs, by behavioral state, (SFig1E-H; see Methods).

### VIP interneuron loss disrupts visual response tuning

Previous work has found that VIP-INs are selectively activated by small visual stimuli (Mesik et al., 2015; Dipoppa et al., 2018) (but see Millman et al., 2020), suggesting that their impact on the surrounding local circuit may depend on visual context. We therefore tested the impact of VIP-IN ablation on the tuning of SST-INs and PNs for drifting grating stimuli of varying sizes (Fig. 2A-B). In control animals, SST-INs exhibited broad tuning for stimulus size, with a preference for stimuli of 20-30° in diameter (Fig. 2A) (Adesnik et al., 2012; Dipoppa et al., 2018). In contrast, PNs were selective for stimuli of ∼10° in diameter and exhibited robust surround suppression that is thought to be mediated in part by SST-IN inhibition (Adesnik et al., 2012) (Fig. 2B). In SST-INs, VIP ablation led to a reduction in the number of visually tuned cells (SFig. 2A-B), decreased size tuning (Fig. 2A,C), and a loss of surround suppression during quiescence (Fig. 2D). In PNs, VIP ablation likewise led to reduced surround suppression across states (Fig. 2B, E-F, SFig. 2 C-D). These changes were associated with a significant increase in the size of preferred stimuli for both SST-INs and PNs (SFig. 2E-H). Together, these results suggest that VIP interneuron activity in V1 normally serves to shape surround suppression and enhance selectivity for small stimuli.

**Figure 2.**
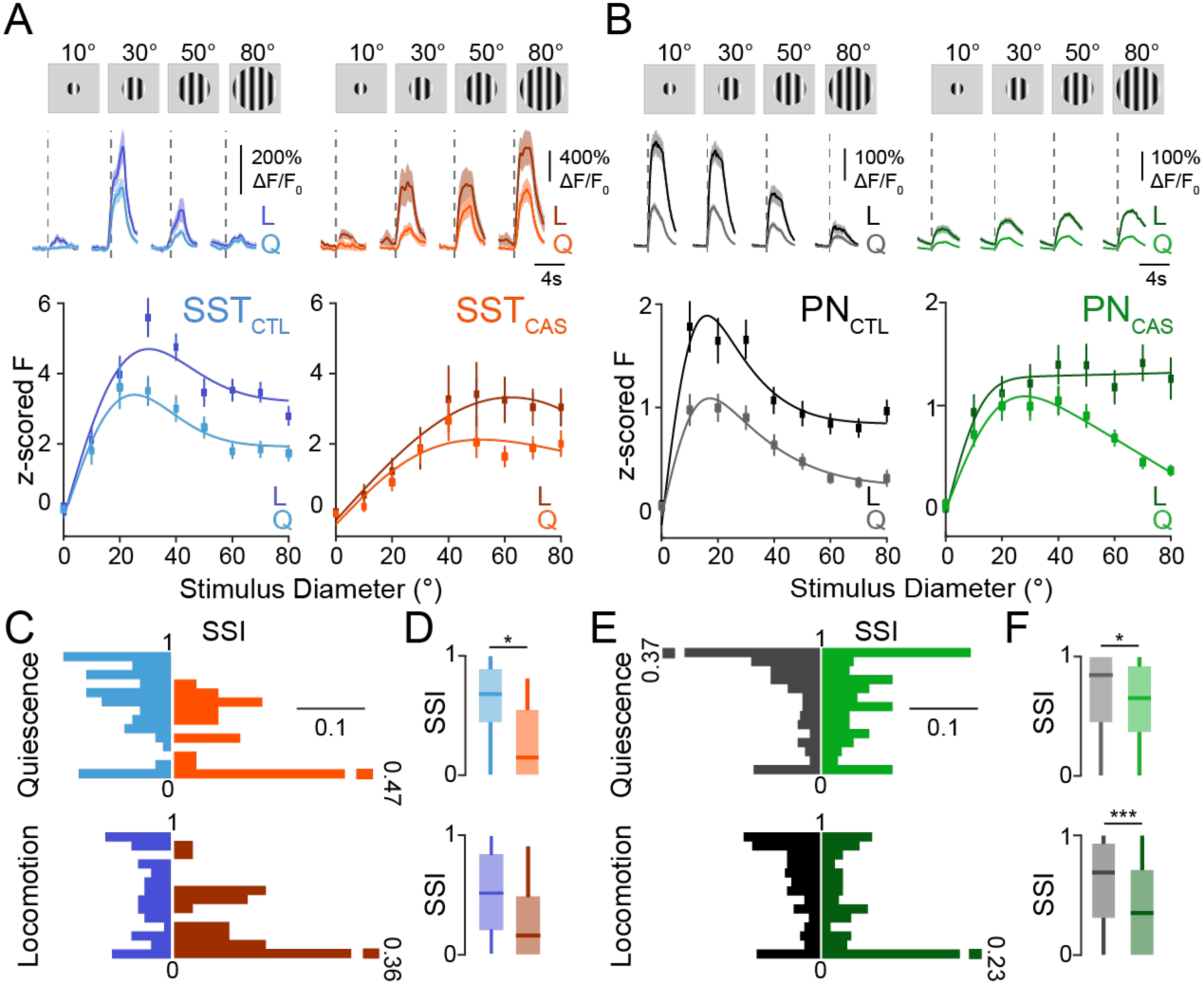
VIP interneuron ablation alters the size tuning properties of SST-INs and PNs. (A) Upper: Responses of example SST-INs to drifting grating stimuli of varying sizes in a control (left, blue; SST_CTL_) and a VIP ablation animal (right, orange; SST_CAS_). Vertical dashed lines indicate visual stimulus onset. Responses during quiescence (Q, light traces) are shown separately from those during locomotion (L, dark traces). Shaded areas indicate mean ± SEM. Lower: Visual responses of SST cells z-scored to the 1s baseline period before the stimulus onset for periods of quiescence (Q, light lines) and locomotion (L, dark lines) for control (blue) and VIP ablation animals (orange). (B) Same as in A, but for PNs in control (gray; PN_CTL_) and VIP ablation (green; PN_CAS_) animals. (C) Probability distribution of the surround suppression index (SSI), separated by locomotion state, for SST-INs in control (blue; Quiescence (upper): n = 86 cells, 6 mice; Locomotion (lower): n = 101 cells, 6 mice) and VIP-ablation animals (orange; Quiescence (upper): n = 36 cells, 4 mice; Locomotion (lower): n = 30 cells, 4 mice). (D) Boxplot of the SSI during quiescence (upper) and locomotion (lower). (E-F) Same as in C-D but for PNs in control (gray; Quiescence (upper): n = 279 cells, 6 mice; Locomotion (lower): n = 314 cells, 6 mice) and VIP ablation (green; Quiescence (upper): n = 175 cells, 5 mice; Locomotion (lower): n = 165 cells, 5 mice) animals. *p<0.05, ***p<0.001, 0/1 inflated beta mixed effects regression model, with experiment type (control or VIP ablation) as fixed effect, mouse with nested imaging field of view as random effect.

### Perturbation of VIP interneurons changes visual perceptual performance

Our findings suggest that VIP-INs function to shape visual responses of interneurons and pyramidal neurons in the local V1 circuit. To examine how VIP-IN regulation of visual feature selectivity may contribute to perceptual behavior, we manipulated VIP-IN activity in V1 during performance of a visual detection task in which head-fixed mice lick for water rewards in response to uncued presentations of contrast-varying stimuli (Fig. 3A- B). Because the extended training times required for expert task performance (see Methods) might permit the emergence of compensatory changes following long-term loss of VIP-INs, we assessed the impact of both short-term (Fig. 3, SFig. 3A-K) and long-term (SFig. 3L-P) manipulation of VIP-IN activity in V1.

**Figure 3.**
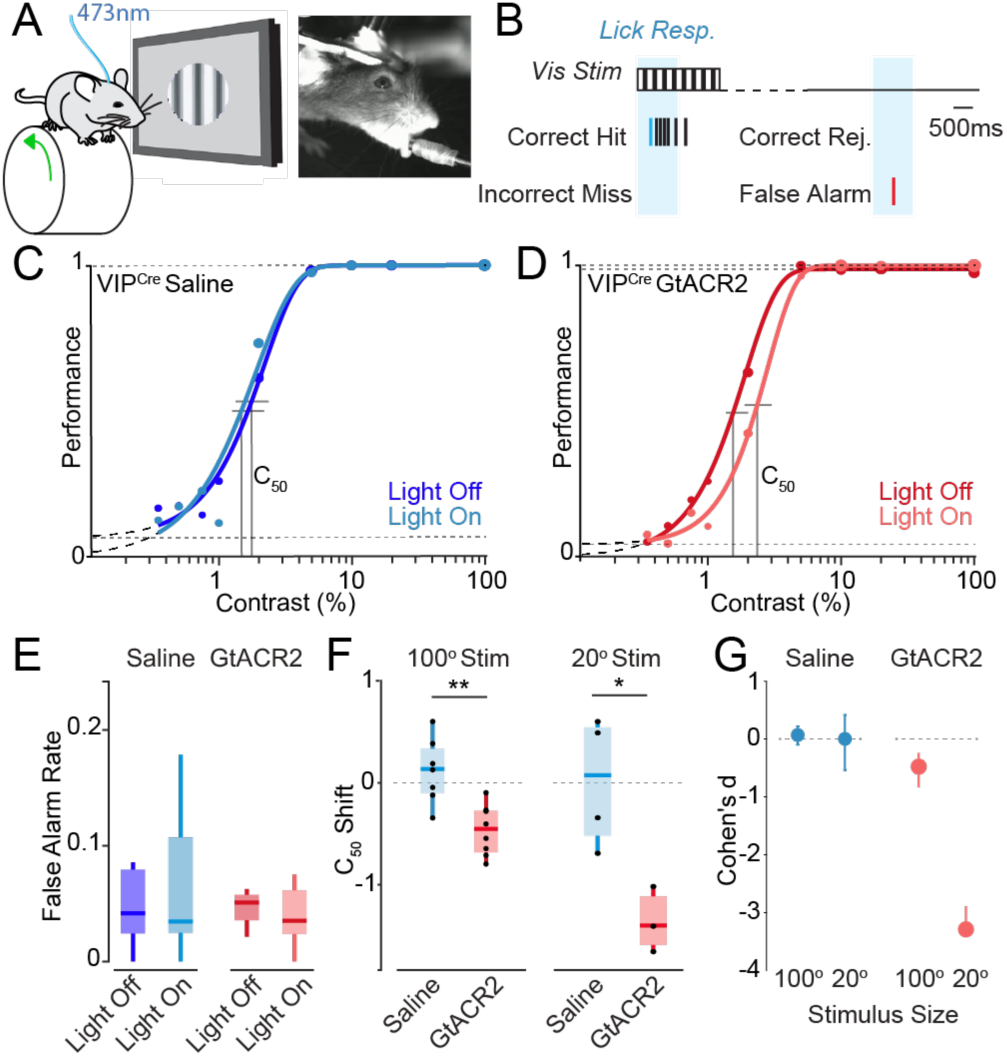
VIP-INs regulate visual perception in a size-dependent manner. (A) Schematic of experimental paradigm with a freely-running head-fixed mouse, lick spout, visual stimulation, and 473 nm optogenetic stimulation. (B) Schematic of visual detection task. (C) Psychometric responses for a representative mouse injected with saline. Darker shade indicates control trials, lighter shade indicates trials with light pulse. The C_50_ is represented by vertical lines. (D) Psychometric responses for a representative mouse injected with the GtACR2 opsin for VIP inhibition upon light stimulation. (E) False alarm rates for control (saline) and GtACR2 mice. (F) C_50_ shift (light off C_50_ – light on C_50_) for control and GtACR2 animals for small (20°) and large (100°) diameter stimuli. (G) Cohen’s d effect size for small (20°) and large (100°) stimuli. 100° stimulus experiments: n = 7 control, 8 GtACR2 mice. 20° stimulus experiments: n = 4 control, 3 GtACR2 mice. *p<0.05, **p<0.01, Student’s t-test.

We expressed a conditional viral construct carrying the light-activated suppressive opsin GtACR2 (Mahn et al., 2018) selectively in either PNs or VIP-INs in V1. Light stimulation to suppress target populations was calibrated using expression of GtACR2 in PNs (SFig. 3A-F). Light stimulation in control animals injected with saline had no impact on performance of the contrast detection task (Fig. 3C). Cell type-specific suppression of VIP-IN activity by activation of GtACR2 with 473nm light caused a rightward shift of the psychophysical performance curve on the task, leading to an increase in the contrast detection threshold (C_50_; Fig. 3D). Ablation of VIP-INs likewise caused a rightward shift in the psychometric performance curve and an overall increase in the C_50_ (SFig. 3L-P) that was particularly prominent during periods of quiescence (SFig. 3O). Neither short nor long-term manipulations of VIP-INs altered false alarm rates or running behavior (Fig.3E, SFig. 3N,P).

Because ablation of VIP-INs had a substantial impact on the size tuning of SST-INs and PNs, we hypothesized that the impact of VIP-IN manipulation on visual perceptual performance may depend on stimulus context. Consistent with our size tuning results, optogenetic suppression of VIP-INs had a substantially larger impact on the detection threshold for small than large stimuli (Fig. 3 F-G, SFig. 3J,K). Overall, these results indicate a role for VIP-INs in regulating the visual feature selectivity of downstream SST-INs and PNs and ultimately in regulating perceptual thresholds for visual stimuli in a size-dependent manner.

## Discussion

Our results reveal a key role for VIP-INs in regulating cortical visual feature selectivity. We find that VIP- IN ablation causes dysregulation of state-dependent modulation of spontaneous activity in SST-INs, but not PNs. We also find that loss of VIP-INs leads to reduced selectivity for visual stimulus size in SST-INs and PNs. We further find that loss or suppression of VIP-IN activity increases the perceptual threshold for detection of visual stimuli. The behavioral impact of VIP-IN manipulation is substantially greater for small than large stimuli, suggesting that the role of VIP-INs in the local V1 circuit varies with visual context.

Imaging in the primary visual cortex of awake control animals revealed robust state-dependent modulation of the activity of both SST-INs and PNs. Several previous studies have found that VIP-INs are activated at the onset of locomotion (Fu et al., 2014; Pakan et al., 2016; Dipoppa et al., 2018) and in response to punishment or unexpected stimuli (Pi et al., 2013; Szadai et al., 2022). VIP-INs receive direct innervation from basal forebrain cholinergic projection neurons (Fu et al., 2014) and are depolarized by application of acetylcholine (Porter et al., 1999; Fu et al., 2014; Askew et al., 2019; Gasselin et al., 2021; Ren et al., 2022), suggesting that they may serve as one avenue by which state information reaches primary cortical circuits. These findings gave rise to the ‘disinhibition model’, where the state-dependent activation of VIP-INs leads to disinhibition of PNs during arousal and locomotion by suppressing intermediary SST-INs (Lee et al., 2013; Fu et al., 2014). Consistent with this view, we found that ablation of VIP-INs caused an enhancement of locomotion modulation of spontaneous activity in local SST-INs, suggesting that VIP-INs normally regulate the degree to which SST-INs are activated during arousal even in the absence of strong visual drive (Millman et al., 2020).

Despite this increase in SST-IN activity, loss of VIP-INs did not change the state modulation of the PN population. Hippocampal VIP-INs comprise several functionally distinct subpopulations, including calretinin-expressing cells that selectively target SST-INs and cholecystokinin-expressing (CCK) cells that directly inhibit the dendrites of PNs (Acsady et al., 1996a; Acsady et al., 1996b; Tyan et al., 2014). Although these subpopulations remain to be fully investigated in the neocortex, anatomical and ex vivo synaptic physiology data suggest that VIP-INs synapse on both SST-INs and on the dendrites of local PNs (Garcia-Junco-Clemente et al., 2017; Chiu et al., 2018). Combined loss of both direct inhibition of PNs and disinhibition via SST-INs following VIP-IN ablation may thus lead to minimal change in PN activity. Alternatively, it is possible that loss of VIP-INs leads to compensatory changes selectively in PNs.

In previous work, we found that developmental perturbation of VIP-INs, which caused a loss of VIP-to SST inhibitory synapses, likewise enhanced state-dependent modulation in SST-INs (Batista-Brito et al., 2017). Developmental perturbation of VIP-INs further abolished state-dependent modulation and substantially impaired visual responses in PNs (Batista-Brito et al., 2017; Mossner et al., 2020). In contrast, adult VIP-IN ablation had no impact on PN state modulation but did affect feature selectivity. Together, these results suggest distinct roles for VIP-INs in the developing and mature cortex.

Cortical networks exhibit dynamic fluctuations across a range of spatial and temporal scales (Cohen and Kohn, 2011). Slow fluctuations, often measured as noise correlations, are a measure of shared variability and can provide insight into functional connectivity (Vinck et al., 2015; Lur et al., 2016), information encoding, and transmission (Cohen and Kohn, 2011; Doiron et al., 2016). Previous work has suggested that inhibition controls the degree to which neural variability is correlated across a cortical population (Stringer et al., 2016). Estimating noise correlations from calcium imaging data presents several challenges. Calcium indicators introduce low pass filtering of neural spiking activity, introducing biases that can be ameliorated by deconvolution (Sabatini, 2019). However, inferring spikes from calcium transients is complex (Pachitariu et al., 2018), partially due to low sensitivity to single action potential events (Huang et al., 2021), and the spike-to-fluorescence transform can vary across populations (Wei et al., 2020). Furthermore, correlations can be biased by comparing samples with mismatched event rates (Cohen and Kohn, 2011). To partially account for these factors, we estimated noise correlations from deconvolved data matched for mean activity levels. We found that loss of VIP-INs enhanced the state-dependent modulation of noise correlations within the SST-IN population, suggesting that inhibition from VIP-INs normally serves to regulate noise correlations of other interneurons. In contrast, we found no consistent impact on PN noise correlations.

Unlike the differential impact of VIP-IN ablation on state-dependent spontaneous SST-IN and PN activity, loss of VIP-INs led to similar changes in the visual responses of both populations. Both SST-INs and PNs exhibited loss of surround suppression and PNs showed an increase in preferred stimulus size following VIP-IN ablation. Previous work has found that SST-INs, which have large receptive fields and receive extensive horizontal cortico-cortical synaptic input, contribute to surround suppression in PNs (Adesnik et al., 2012). VIP- IN inhibition of SST-INs regulates the sensitivity of PNs to visual stimulus features in the surround (Keller et al., 2020). We found that, despite their small receptive fields (Mesik et al., 2015; Dipoppa et al., 2018), VIP-INs contribute to selectivity for stimulus size in SST-INs, suggesting that inhibitory synaptic interactions may likewise mediate surround suppression in interneuron populations. The loss of tuning and shift toward decreased surround suppression in SST-INs following VIP-IN ablation was associated with a similar shift and increased preferred stimulus size in local PNs. The effects of VIP-IN ablation on size tuning were observed across quiescent and aroused behavioral states, suggesting that the impact of VIP-INs on visual tuning may be independent from their role in state-dependent regulation of spontaneous activity. Overall, our results suggest that VIP-INs robustly regulate size tuning in the V1 circuit. However, the synaptic mechanisms by which they promote tuning for smaller stimuli during visually driven activity patterns remain to be further explored.

In good agreement with previous work (Cone et al., 2019), we found that suppression of VIP-IN activity in V1 led to an increased perceptual threshold for visual contrast detection in behaving mice. Both short and long-term suppression of VIP-IN activity increased perceptual thresholds without affecting false alarm rates or locomotion. However, the behavioral impact of VIP-IN suppression was greatly enhanced for small compared to large visual stimuli, suggesting that reduced PN size tuning in the absence of VIP-IN activity selectively impairs the animal’s ability to detect small stimuli. Together, our results indicate that VIP-INs play a substantial role in regulating cortical feature selectivity and that their impact on sensory processing varies with visual context.

**Table 1.** Summary table of all statistical tests.

| Figure | Comparison |  | N | Test |  | Statistics |  |  |  |
| --- | --- | --- | --- | --- | --- | --- | --- | --- | --- |
|  |  |  |  |  |  | Estimate | t-stat | CI | P-value |
| 1F | Comparison of locomotion modulation indices for SST control and SST with VIP ablated | | Ctrl: 7 mice, 600 cells<br>w/out VIP: 5 mice, 277 cells | Linear mixed effects model<br>$y \sim \text{experimentType} + (1 \mid \text{mouse:Fov})$ | | 0.140 | 2.566 | [0.033, 0.247] | **0.010** |
| 1I | Comparison of locomotion modulation indices for PN control and PN with VIP ablated | | Ctrl: 6 mice, 1687 cells<br>w/out VIP: 6 mice, 1617 cells | Linear mixed effects model<br>$y \sim \text{experimentType} + (1 \mid \text{mouse:Fov})$ | | 0.050 | 1.817 | [-0.004, 0.104] | 0.067 |
| 2D | Comparison of surround suppression indices for SST control and SST with VIP ablated | Q | Ctrl: 6 mice, 86 cells<br>w/out VIP: 4 mice, 30 cells | Zero/one inflated beta mixed effects regression model<br>(Experiment type as fixed effect; mouse with nested FoV as random effects) |  | -0.930 | -3.620 | [-1.439, -0.422] | **p<0.001** |
|  |  | L | Ctrl: 6 mice, 101 cells<br>w/out VIP: 4 mice, 36 cells |  |  | -0.217 | -0.909 | [-0.684, 0.250] | 0.366 |
| 2F | Comparison of surround suppression indices for PN control and SST with VIP ablated | Q | Ctrl: 6 mice, 279 cells<br>w/out VIP: 5 mice, 175 cells | Zero/one inflated beta mixed effects regression model<br>(Experiment type as fixed effect; mouse with nested FoV as random effects) |  | -0.545 | -4.288 | [-0.795, -0.295] | **p<0.001** |
|  |  | L | Ctrl: 6 mice, 314 cells<br>w/out VIP: 5 mice, 165 cells |  |  | -0.423 | -2.775 | [-0.724, -0.123] | **0.006** |
| 3E | Comparison of false alarm rates for light stimulation on trials and light off trials | Saline | 7 mice | Paired t-test |  |  |  | [-0.063, 0.052] | 0.812 |
|  |  | GtACR2 | 8 mice |  |  |  |  | [-0.019, 0.040] | 0.444 |
| 3F | Comparison of C50 shift for saline injected and GtACR2 injected (opto) mice | 100° stim | Saline: 7 mice; GtACR2: 8 mice | Unpaired t-test |  |  |  | [0.268, 0.897] | **0.002** |
|  |  | 20° stim | Saline: 4 mice; GtACR2: 3 mice |  |  |  |  | [0.338, 2.459] | **0.019** |
| S1C | Comparison of VIP cell density in control and VIP ablated mice | 10 days post-inject | Saline: 3 mice; Caspase: 3 mice | Unpaired t-test |  |  |  | [-51.53, 175.52] | 0.204 |
|  |  | 14 days post-inject | Saline: 3 mice; Caspase: 3 mice |  |  |  |  | [76.07, 283.96] | **0.009** |
|  |  | 21 days post-inject | Saline: 3 mice; Caspase: 3 mice |  |  |  |  | [148.66, 275.64] | **p < 0.001** |
| S1D | Comparison of VIP cell density in behavioral control mice and behavioral VIP ablated mice |  | Saline: 4 mice; Caspase: 4 mice | Unpaired t-test |  |  |  | [151.21, 309.09] | **p < 0.001** |
| S1E | Noise correlations of $\Delta F/F_0$ | SST | Ctrl: 6 mice, 2180 cell pairs<br>w/out VIP: 4 mice, 447 cell pairs | Linear mixed effects model<br>$y \sim 1 + \text{state} * \text{experimentType} + (1 \mid \text{mouse:Fov})$ | State | 0.045 | 4.282 | [0.024, 0.065] | **p < 0.001** |
|  |  |  |  |  | ExpType | -0.031 | -0.580 | [-0.135, 0.073] | 0.562 |
|  |  |  |  |  | State * ExpType | 0.071 | 2.812 | [0.022, 0.121] | **0.005** |
| | | PN | Ctrl: 5 mice, 1077 cell pairs<br>w/out VIP: 6 mice, 2481 cell pairs | Linear mixed effects model<br>$y \sim 1 + \text{state} * \text{experimentType} + (1 \mid \text{mouse:Fov})$ | State | 0.007 | 0.569 | [-0.016, 0.029] | 0.569 |
|  |  |  |  |  | ExpType | 0.023 | 1.130 | [-0.017, 0.064] | 0.259 |
|  |  |  |  |  | State * ExpType | 0.024 | 1.763 | [-0.003, 0.051] | 0.078 |
| S1F | Noise correlations of deconvolved $\Delta F/F_0$ | SST | Ctrl: 6 mice, 2180 pairs<br>w/out VIP: 4 mice, 447 pairs | Linear mixed effects model<br>$y \sim 1 + \text{state} * \text{experimentType} + (1 \mid \text{mouse:Fov})$ | State | -0.011 | -1.424 | [-0.026, 0.004] | 0.155 |
|  |  |  |  |  | ExpType | -0.027 | -0.640 | [-0.112, 0.057] | 0.522 |
|  |  |  |  |  | State * ExpType | 0.099 | 5.322 | [0.062, 0.135] | **p < 0.001** |
| | | PN | Ctrl: 5 mice, 1077 pairs<br>w/out VIP: 6 mice, 2481 pairs | Linear mixed effects model<br>$y \sim 1 + \text{state} * \text{experimentType} + (1 \mid \text{mouse:Fov})$ | State | -0.016 | -1.671 | [-0.035, 0.003] | 0.095 |
|  |  |  |  |  | ExpType | -0.019 | -2.337 | [-0.035, -0.003] | 0.020 |
|  |  |  |  |  | State * ExpType | 0.015 | 1.336 | [-0.007, 0.038] | 0.181 |
| S1G/H | Noise correlations of mean-matched deconvolved $\Delta F/F_0$ | SST | Ctrl: 6 mice, (1293, 975) (Q, L) pairs<br>w/out VIP: 4 mice, (261, 202) (Q, L) pairs | Linear mixed effects model<br>$y \sim 1 + \text{state} * \text{experimentType} + (1 \mid \text{mouse:Fov})$ | State | -0.024 | -1.476 | [-0.056, 0.008] | 0.140 |
|  |  |  |  |  | ExpType | -0.014 | -0.262 | [-0.120, 0.092] | 0.793 |
|  |  |  |  |  | State * ExpType | 0.098 | 3.176 | [0.037, 0.158] | **0.0015** |
|  |  | PN Control | Ctrl: 6 mice, (748, 748) (Q, L) pairs | Linear mixed effects model | State | -0.017 | -1.523 | [-0.039, 0.005] | 0.128 |
|  |  |  | w/out VIP: 6 mice, (1658, 1786) (Q, L) pairs | y ~ 1 + state*experimentType + (1 \| mouse:FoV) | ExpType | -0.015 | -1.508 | [-0.033, 0.004] | 0.132 |
|  |  |  |  |  | State * ExpType | 0.011 | 0.846 | [-0.015, 0.038] | 0.398 |
| S2B | Comparison of percent visually tuned cells of all visually responsive cells in SST controls and SST with VIP ablated |  | Ctrl: 6 mice<br>w/out VIP: 4 mice | Unpaired t-test |  |  | 2.844 | [0.060, 0.577] | **0.022** |
| S2D | Comparison of percent visually tuned cells of all visually responsive cells in PN controls and PN with VIP ablated |  | Ctrl: 6 mice<br>w/out VIP: 5 mice | Unpaired t-test |  |  | 1.582 | [-0.052, 0.296] | 0.148 |
| S2F | Comparison of preferred size of visual responses in SST controls and SST with VIP ablated | Q | Ctrl: 6 mice, 86 cells<br>w/out VIP: 4 mice, 30 cells | Zero/one inflated beta mixed effects regression model (Experiment type as fixed effect; mouse with nested FoV as random effects) |  | 43.12 | 2.971 | [14.67, 71.57] | **0.004** |
|  |  | L | Ctrl: 4 mice, 66 cells<br>w/out VIP: 4 mice, 21 cells |  |  | 10.74 | 0.605 | [-24.06, 45.54] | 0.547 |
| S2H | Comparison of preferred size of visual responses in PN controls and PN with VIP ablated | Q | Ctrl: 6 mice, 254 cells,<br>w/out VIP: 5 mice, 175 cells | Zero/one inflated beta mixed effects regression model (Experiment type as fixed effect; mouse with nested FoV as random effects) |  | 22.70 | 3.040 | [8.07, 38.33] | **0.003** |
|  |  | L | Ctrl: 3 mice, 202 cells<br>w/out VIP: 3 mice, 122 cells |  |  | 1.89 | 0.201 | [-16.49, 20.27] | 0.841 |
| S3F | Comparison of modulation indices in response to various light intensities | Low vs Medium | 1 mouse, 84 units | Friedman's ANOVA and post-hoc Wilcoxon signed rank test |  |  |  |  | **p < 0.001** |
|  |  | Medium vs High |  |  |  |  |  |  | **p < 0.001** |
|  |  | Low vs High |  |  |  |  |  |  | **p < 0.001** |
| S3H | Comparison of run probability in saline injected and GtACR2 injected mice |  | Saline: 7 mice<br>GtACR2: 8 mice | Linear mixed effects model y ~ experimentType + (1 \| mouse:FoV) |  | 0.068 | 0.742 | [-0.114, 0.250] | 0.460 |
| S3I | Comparison of c50 shift for saline injected and GtACR2 injected (opto) mice, separated by behavioral state | Q | Saline: 7 mice<br>GtACR2: 8 mice | Unpaired t-test |  |  | 3.615 | [0.276, 1.094] | **0.003** |
|  |  | L |  |  |  |  | 1.263 | [-0.239, 0.913] | 0.229 |
| S3J | Comparison of c50 for light stimulation on trials and light off trials in GtACR2 mice | 100° stim | GtACR2: 8 mice | Paired t-test |  |  | -5.394 | [-0.675, -0.263] | **0.001** |
|  |  | 20° stim | GtACR2: 3 mice | Paired t-test |  |  | -7.273 | [-2.189, -0.562] | **0.018** |
| S3K | Comparison of c50 for light stimulation on trials and light off trials in Saline mice | 100° stim | Saline: 7 mice | Paired t-test |  |  | 0.944 | [-0.181, 0.407] | 0.382 |
|  |  | 20° stim | Saline: 4 mice | Paired t-test |  |  | 0.072 | [-1.001, 1.048] | 0.947 |
| S3N | Comparison of run probability in saline injected and VIP-ablated mice |  | Ctrl (saline): 14 mice<br>w/out VIP: 8 mice | Linear mixed effects model y ~ experimentType + (1 \| mouse:FoV) |  | 0.0319 | 0.729 | [-0.054, 0.118] | 0.467 |
| S3O | Comparison of c50 for saline injected and VIP-ablated mice, separated by behavioral state | Q | Ctrl (saline): 14 mice<br>w/out VIP: 8 mice | Unpaired t-test |  |  | -2.231 | [-1.655, -0.056] | **0.037** |
|  |  | L |  |  |  |  | -1.403 | [-2.054, 0.402] | 0.176 |
| S3P | Comparison of lick probability for lowest contrast stimuli in saline injected and VIP-ablated mice |  | Ctrl (saline): 14 mice<br>w/out VIP: 8 mice | Unpaired t-test |  |  | 1.663 | [-0.010, 0.095] | 0.109 |

## Acknowledgements

The authors thank all members of the Higley and Cardin laboratories for helpful input throughout all stages of this study. We thank Rima Pant for generation of AAV vectors and Quentin Perrenoud for guidance on the behavioral task design. This work was supported by funding from the NIH (R01EY022951 R01MH113852 to JAC, K99EY030549 to KAF, EY026878 to the Yale Vision Core), an award from the Kavli Institute of Neuroscience (to JAC), support from the Ludwig Foundation (to JAC), and a BBRF Young Investigator Grant (to KAF).

## Author Contributions

KAF, SL, and JAC designed the experiments. KAF, JS, CA, and HS collected the data. KAF analyzed the data. KAF and JAC wrote the manuscript.

## Conflicts of Interest

The authors declare no conflicts of interest exist.

## Materials and Methods

### Animals

All animal handling and maintenance was performed according to the regulations of the Institutional Animal Care and Use Committee of the Yale University School of Medicine. Adult male and female C57BL/6J VIP-IRES-Cre^+/0^ (Jax stock no. 031628), Emx1-IRES-Cre^+/0^ (Jax stock no. 005628), VIP-IRES-Cre^+/0^ mice crossed with Sst-IRES-Flp^+/0^ (Jax stock no. 031629) mice, VIP-IRES-Cre^+/0^ mice crossed with Ai9^F/0^ (Jax stock no. 007909), VIP-IRES-Cre^+/0^ crossed with Sst-IRES-Flp^+/0^, and Sst-IRES-Cre^+/0^ (Jax stock no. 018973) crossed with Ai148^F/0^ (Ai148(TIT2L-GC6f-ICL-tTA2)-D, Jax stock no. 030328) mice were kept on a 12h light/dark cycle, provided with food and water ad libitum, and housed individually following headpost implants. Imaging experiments were performed during the light phase of the cycle.

### Surgical Procedures

Surgeries were performed in adult mice (P90–P180) in a stereotaxic frame, anesthetized with 1-2% isoflurane mixed with pure oxygen. Injections were made via beveled glass micropipette at a rate of 40-60 nl/min into the primary visual cortex (V1) at a depth of L2/3 (∼350 um) (QSI, Stoelting Co.). For imaging experiments, we injected 200nL of adenoassociated virus (AAVdj-ef1a-fDIO-GCaMP6m (plasmid gift of K. Deisseroth lab, Stanford), AAV5-Syn-GCaMP6s (Addgene # 100843), or AAV5-Syn-FLEX-GCaMP6s (Addgene # 100845); diluted to a titer of 10^12) unilaterally at three sites (in mm from Bregma): AP 3.5, ML 1.5, DV 0.4; AP 3, ML 2, DV 0.4; AP 2.5, ML 2.5, DV 0.4. We also injected 1 uL of either the Cre-dependent Caspase-3 virus (AAV5 ef1α-Flex-taCasP3-TEVP; UNC Vector Core) or saline into V1 (in left V1 for imaging experiments, and bilaterally for behavioral experiments, 2-2.5 mm lateral and 3.5-4.0 mm posterior from Bregma). For behavioral optogenetic experiments, we bilaterally (2-2.5 mm lateral and 3.5-4.0 mm posterior from Bregma) injected 1 uL of the conditional GtACR2 virus (AAV5-hSyn1-SIO-stGtACR2-FusionRed, Addgene #105677), or 1uL saline injection for controls. After injection, pipettes were left in the brain for 5-10 min to prevent backflow.

For headpost implantation, mice were anesthetized with isoflurane and the scalp was cleaned with Betadine solution. An incision was made at the midline and the scalp resected to each side to leave an open area of the skull. After cleaning the skull and scoring it lightly with a surgical blade, a custom titanium head post was secured with C&B-Metabond (Butler Schein) with the left V1 centered. Two skull screws (McMaster-Carr) were placed at the right anterior and posterior poles (bilateral to the injection site). A 3 mm^2^ craniotomy was made over the left V1. A glass window made of a 3 mm^2^ rectangular inner cover slip adhered with an ultraviolet curing adhesive (Norland Products) to a 5 mm round outer cover slip (both #1, Warner Instruments) was inserted into the craniotomy and secured to the skull with Cyanoacrylate glue (Loctite). A circular ring was attached to the titanium headpost with glue, and additional Metabond was applied to cover any exposed skull and to cover each skull screw. For the behavioral experiments, two skull screws (McMaster-Carr) were placed at the right anterior and posterior poles (bilateral to the injections/cranial window). Two nuts (McMaster-Carr) were glued in place over the bregma point with cyanoacrylate and secured with C&B-Metabond (Butler Schein). The Metabond was extended along the sides and back of the skull to cover each screw. For optogenetics experiments, two fiber-optic cannulas were lowered directly over the bilateral virus injection sites and secured with dental cement to allow for the delivery of light directly onto the surface of the cortex during the behavioral task. Analgesics were given immediately after surgery (5 mg/kg Carprofen and 0.05 mg/kg Buprenorphine) and on the two following days to aid recovery. Mice were given a course of antibiotics (Sulfatrim, Butler Schein) to prevent infection and were allowed to recover for 3-5 days following implant surgery before beginning wheel training.

### Histology

Following experiments, animals were given a lethal dose of sodium pentobarbital and perfused intracardially with 0.9% saline followed by cold 4% paraformaldehyde in 0.1 m sodium phosphate buffer. For the Caspase-3 virus efficacy and timeline experiments, VIP-Cre^+/^0;Ai9F^/0^ animals were perfused 10, 14, and 21 days after unilateral injection of the AAV-ef1α-Flex-taCasP3-TEVP virus. Brains were removed and fixed in 4% PFA/PBS solution for 24 hours and subsequently stored in PBS. Tissue was sectioned at 40μm using a vibrating blade microtome, mounted, and visualized by light microscopy. Widefield images were acquired on an Olympus BX53 fluorescence microscope. In a subset of cases, confocal images were taken with a Zeiss LSM 900.

To minimize counting bias we compared sections of equivalent bregma positions, defined according to the Mouse Brain atlas (Franklin and Paxinos, 2013). The total number of cells expressing tdTomato (from the Ai9 reporter mouse line) were counted for a defined optical area within V1. Cell counting was performed manually using a standardized 100 um x 100 um grid overlay to determine the average VIP cell density in layers 2/3 of V1 across three consecutive sections. The percentages of VIP interneurons were calculated as a ratio between the total number of tdTomato^+^ cells in the injected area over the total number of tdTomato^+^ cells on the contralateral control side.

### In Vivo Calcium Imaging

All imaging was performed during the second half of the light cycle in awake, behaving mice that were head-fixed so that they could freely run on a cylindrical wheel (Vinck et al., 2015; Batista-Brito et al., 2017; Tang and Higley, 2020). A magnetic angle sensor (Digikey) attached to the wheel continuously monitored wheel motion. Mice received at least three wheel-training habituation sessions before imaging to ensure consistent running bouts. The face (including the pupil and whiskers) was imaged with a miniature CMOS camera (Blackfly s-USB3, Flir) with a frame rate of 10 Hz.

Imaging was performed using a resonant scanner-based two-photon microscope (MOM, Sutter Instruments) coupled to a Ti:Sapphire laser (MaiTai DeepSee, Spectra Physics) tuned to 920 nm for GCaMP6. Emitted light was collected using a 25x 1.05 NA objective (Olympus). Mice were placed on the wheel and head fixed under the microscope objective. To prevent light contamination from the display monitor, the microscope was enclosed in blackout material that extended to the headpost. Images were acquired using ScanImage 4.2 at 30 Hz, 512x512 pixels. Imaging of layer 2/3 was performed at 150-350 μm depth relative to the brain surface. For each mouse, 1-4 fields of view were imaged. Visual stimulation, wheel position, and Ca2+ imaging microscope resonant scanner frame ticks, were digitized (5 kHz) and collected through a Power 1401 (CED) acquisition board using Spike 2 software. During each session, spontaneous activity was collected for 10 mins before the series of visual stimuli were presented, and 10 mins after (20 mins total) as the mouse freely moved on the wheel in front of a mean-luminance gray screen.

### Visual Stimulation

Visual stimuli were generated using Psychtoolbox-3 in MATLAB and presented on a gamma-calibrated LCD monitor (17 inches) at a spatial resolution of 1280 x 960, a real-time frame rate of 60Hz, and a mean luminance of 30 cd/m^2^ positioned 20 cm from the right eye. Stimuli had a temporal frequency of 2 Hz, spatial frequency of 0.04 cycles per degree, and orientation of 180°. To center stimuli on the receptive field, 100% contrast stimuli were randomly presented in nine 3x3 sub-regions to identify the location that evoked the largest population response in the field of view. The screen was centered, and the process was repeated until a center was identified. Stimuli in each session were randomized and presented in blocks with a fixed duration of 2 s and an interstimulus interval of 5 s, with a mean-luminance gray screen between stimuli. For size tuning, the visual angle was linearly spaced from 0 to 80° in diameter in steps of 10°, where each size was presented 45 times.

### Visual Detection Task

Mice were trained to perform a visual contrast detection task while head-fixed on a wheel. Mice were placed on a water-controlled schedule with careful weight monitoring and habituated to head fixation. Once mice were stabilized at 83-86% of their starting weight and exhibited consistent running bouts on the wheel, they were trained to lick a waterspout in response to the presentation of a high-contrast (100%), full-screen stimulus of 1 s duration (temporal frequency: 2Hz; spatial frequency: 0.05 cycles/degree). Successful detection of the visual stimulus resulted in a reward and prolonged the presentation for an additional 1 s (total 2 s). When a performance criterion of > 95% hit rates and < 10% false alarm rates was reached (∼5-10 days), they were moved to a psychometric version of the task where the stimulus contrast varied randomly across trials. Contrast was selected on each trial from the series: 0, 0.35, 0.5, 0.75, 1, 2, 5, 10, 20, 100%. To determine how visual perception is regulated in a size-dependent manner, either small (20° diameter) or large (100° diameter; full screen) stimuli were used throughout the duration of the task. Stimulus and non-stimulus (0% contrast) trials were presented at a one-to-one ratio. To maintain motivation in the task, high contrasts were over sampled such that stimuli greater than 1.5% contrast made up 70% of the displayed (non-zero) stimuli. The response window for a correct response (hit) began at stimulus onset and lasted for 1 second. Hits were rewarded with a small (3 μl) drop of water. The inter-trial interval (ITI) for both stimulus and non-stimulus trials was drawn from an exponential distribution to ensure a flat hazard rate, with a mean ITI of 4 seconds, a minimum ITI of 1.5 seconds, and a maximum ITI of 10 seconds. False alarms were punished with an extended inter-trial interval by re-sampling the ITI starting from the time of the false alarm. Mice performed the task daily for 45 min per session, over 10 sessions. Mice began the contrast detection task no earlier than 22 days following virus injection.

To acutely inhibit VIP-INs, fiber-optic cannulas were surgically implanted at the injection sites, bilaterally in V1. The light was delivered through a fiber coupled 473nm LED laser to the cortical surface at an intensity of 75 mW/mm^2^ (Cardin, 2010). Optogenetic stimulation trials were randomly assigned to 50% of the stimulus and non-stimulus trials, where a 2.25s pulse of light was activated 250ms preceding the onset of the visual stimulus.

### Data analysis

#### Wheel Position and Changepoints

Wheel position was determined from the output of the linear angle detector. The circular wheel position variable was first transformed to the [-π, π] interval. The phases were then circularly unwrapped to get running distance as a linear variable, and locomotion speed was computed as a differential of distance (cm/s). A change point detection algorithm detected locomotion onset/offset times based on changes in standard deviation of speed. Locomotion onset or offset times were estimated from periods when the moving standard deviations, as determined in a 0.5s window, exceeded or fell below an empirical threshold of 0.1. Locomotion trials were required to have average speed exceeding 0.5 cm/s and last longer than 1 s. Quiescence trials were required to last longer than 2 s and have an average speed < 0.5 cm/s.

#### Quantification of Calcium Signals

Analysis of imaging data was performed using ImageJ and custom routines in MATLAB (The Mathworks). Motion artifacts and drifts in the Ca 2+ signal were corrected with the moco plug-in in ImageJ (Dubbs et al., 2016), and regions of interest (ROIs) were selected as previously described (Chen et al., 2013). All pixels in each ROI were averaged as a measure of fluorescence, and the neuropil signal was subtracted (Chen et al, 2013; Lur et al., 2016; Tang et al., 2020). ΔF/F was calculated as (F-F_0_)/F_0_, where F_0_ was the lowest 10% of values from the neuropil-subtracted trace for each session.

#### Modulation Index

For modulation by behavioral state without visual stimulation, we used the spontaneous periods recorded as described above and selected locomotion trials that lasted 5 s or longer and quiescent trials that lasted 30 s or longer. To determine whether Ca 2+ activity was altered during behavioral state transitions, ΔF/F(t) from [0,5]s after locomotion onset (Ca_L-ON_) was compared with ΔF/F(t) from [20,25]s after locomotion offset (Ca_Q_) by computing a modulation index (MI), where MI = (Ca_L-ON_ – Ca_Q_)/(Ca_L-ON_ +Ca_Q_). A minimum of 5s of quiescence after this period [25,30]s was required to prevent anticipatory effects on Ca_Q_. To ascertain the significance of this MI, we used a shuffling method in which the wheel trace was randomly circularly shifted relative to the fluorescence trace 1,000 times. Cells were deemed significantly modulated if their MI was outside of the 95% confidence interval of the shuffled comparison.

#### Visual Responses

Visual response amplitude was calculated as the z-scored change in fluorescence (z-scored F) during the 2s visual stimulus compared to the 1s baseline before the stimulus. To separate effects of state, the mouse was required to be running (or sitting) during the full duration of the 1s baseline and the 2 s visual stimulation. To determine if cells were visually responsive, a bootstrapped null distribution was generated by sampling with replacement from each cell’s pre-stimulus baseline. Cells were deemed visually responsive if their mean responses to their preferred stimulus (100% contrast or preferred stimulus size) was outside of the 95% bootstrapped confidence interval in at least one of the two behavioral state conditions (quiescence or locomotion).

Size tuning of all cell types, and particularly of SST-INs, prefer larger stimulus sizes when not well centered (Dipoppa et al., 2018). After centering our stimuli, we only included cells in our size tuning analysis that were both visually responsive and tuned and thus well matched to the visual stimuli. To identify tuned cells, size tuning curves were fitted by least-squares with the difference in error functions (*erf*):

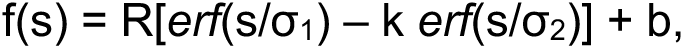

where s is the size of the stimulus, and the free parameters are R, k, b, σ_1_ and σ_2_. Tuned cells were defined as visually responsive cells whose fit curve was not monotonically increasing or decreasing. To compute the suppression index (SI), we normalized the z-scored F of tuned cells to their peak and computed the difference in normalized activity at the preferred size and the largest stimuli (Adesnik et al., 2012). SI index was computed using the preferred size based on stimuli presented, not on the fit data, to prevent it from being affected by goodness of fit.

Pairwise correlations were determined as trial-by-trial fluctuations in response strength between cells to high contrast stimuli (80-100%). To compute these noise correlation coefficients, either the ΔF/F_0_ traces were used, or the spike traces for each cell were deconvolved using a first order autoregressive model (OASIS, (Friedrich et al., 2017)), and spike times were selected as those that exceeded 3 standard deviations. State conditions were separated (v = 0 stationary, v = 1 running). For each trial in state v for each of the [80, 90, 100]% contrast trials (stimulus s) (t_vs_) we calculated the average number of spikes (r) for each cell (i) in cell class I: ^r̅^_ivs_=〈r(t_vs_,i)〉, where t_vs_∈{v,s} and i∈{c}. As differences in firing rates can markedly affected measured noise correlations (Cohen and Kohn, 2011), in a subset of analyses firing rates between stationary and running conditions were mean-matched across cells using a threshold of 0.5 Hz. To do so, for each cell’s locomotion firing rate, we identified a “paired” cell whose quiescence firing rate fell within the threshold. If none existed, we eliminated that cell. For each t_vs_ (trial in locomotion condition v with stimulus s) and for each i (cell), we subtracted the average response, Δr_ivs_ = r(t_vs_,i) - ^r̅^_ivs_, and pooled the trial-by-trial fluctuations across stimuli (Δr_iv_). The Pearson’s correlation of these spike count responses Δr_iv_ across pairs of cells (⍴_ij,v_ for cells i,j during condition v) was calculated.

#### Behavioral Performance

We restricted the data used per session automatically as follows. First, we ensured that when the mice stopped performing at the end of a session, these data were not incorporated into the average. This was done by computing a 10-point moving average of the data. For the k-th trial, we then computed the average performance of the mouse (as hit_O_) until the (k-1)-th trial. This average performance was computed starting from the trial where the mouse had obtained at least 10 rewards, to prevent poor performance at the start from influencing the average. (Note that the first ten trials in each session were 100% contrast trials). The last trial was defined as the trial at which the 10-point moving average of the hit rate (HR) fell below 75% of the mean performance up to that point and did not recover above this level anymore. We then computed the average HR for each contrast, and the average false alarm rate (FAR) from the non-stimulus (0% contrast) trials. In the optogenetic experiments, performance and FAR were further separated by the presence of the light stimulus (“light-off” or “light-on” trials). Sessions were removed from the analysis if the median light-off FAR or HR at the two lowest contrasts (0.35% and 0.5%) exceeded 40% or if the median light-off HR at the highest contrast (100%) was below 75%.

Performance was separated by arousal state, where locomotion states were indicated by any duration of locomotion in the 1s preceding the visual stimulation. As trials for each session were categorized by state and light stimulation (for the optogenetic experiments), and thresholds gleaned from psychometric curve fitting are sensitive to low trial numbers, we aggregated our trials across 10 sessions to optimize our fits. We fit the psychometric curves with a Weibull function using the *psignifit* toolbox in MATLAB, a software package which implements the maximum-likelihood method (Frund et al., 2011). A 95% confidence interval was determined by the percentile bootstrap method implemented by *psignifit* based on 2000 simulations. The contrast detection threshold (C_50_) was the lowest contrast that can be detected at least 50% of the time, scaled by the guess rate (lower asymptote) and lapse rate (upper asymptote) of the psychometric curve fits. We calculated the C_50_ *shift* by subtracting the C_50_ value for light-on trials from the C_50_ value for light-off trials to measure the difference in performance after inhibiting VIP-INs.

#### Statistical Analyses

We used mixed effect regression models for imaging data, due to its nested structure with multiple cells recorded within each mouse. We treated the experiment type (control or VIP-ablated) as the fixed effect, and the individuals (mice) were random effects. Since our experimental design was between-subject, we used the *lmefit* function in MATLAB to fit the intercepts of the random effects, with the response modeled as:

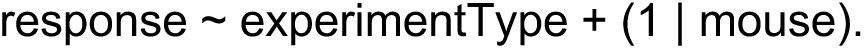

which has the following mathematical form:

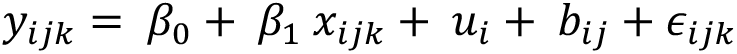

where *y_ijk_* is the *i^th^* observation for the *j^th^* mouse in the *k^th^* field of view. *x_ijk_* is the experiment type (control or VIP-ablated) for the observation *i* of the *j^th^* mouse and *k^th^* field of view. β_0_ is the intercept, β_1_ is the effect of the experiment type, *u_i_* is the random effect for the *i^th^* observation, *b_ij_* is the random effect for the *i^th^* observation in the *j^th^* mouse, and *∈_ijk_* is the residual error for the i^th^ observation within the j^th^ mouse and *k^th^* field of view. The random effects have prior distributions 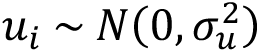 and 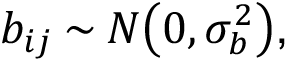, and the error term has the distribution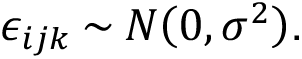.

Since the surround suppression index and the preferred size were continuous bounded variables, we instead used a 0/1 inflated beta mixed effect regression model. For these data, preferred size was scaled to [0, 1] to represent the proportion of the size compared to maximum. Then we fit a 0/1 inflated beta mixed effect regression model using the *gamlss* package in R using the family “BEINF”.

For the behavioral data, given one psychometric curve fit per mouse, data was compared with a t-test after a test for normality. Paired t-tests were used when comparing light-on and light-off trials, and unpaired t-tests were used when comparing across mice.

---

**Supplementary Figure 1.**
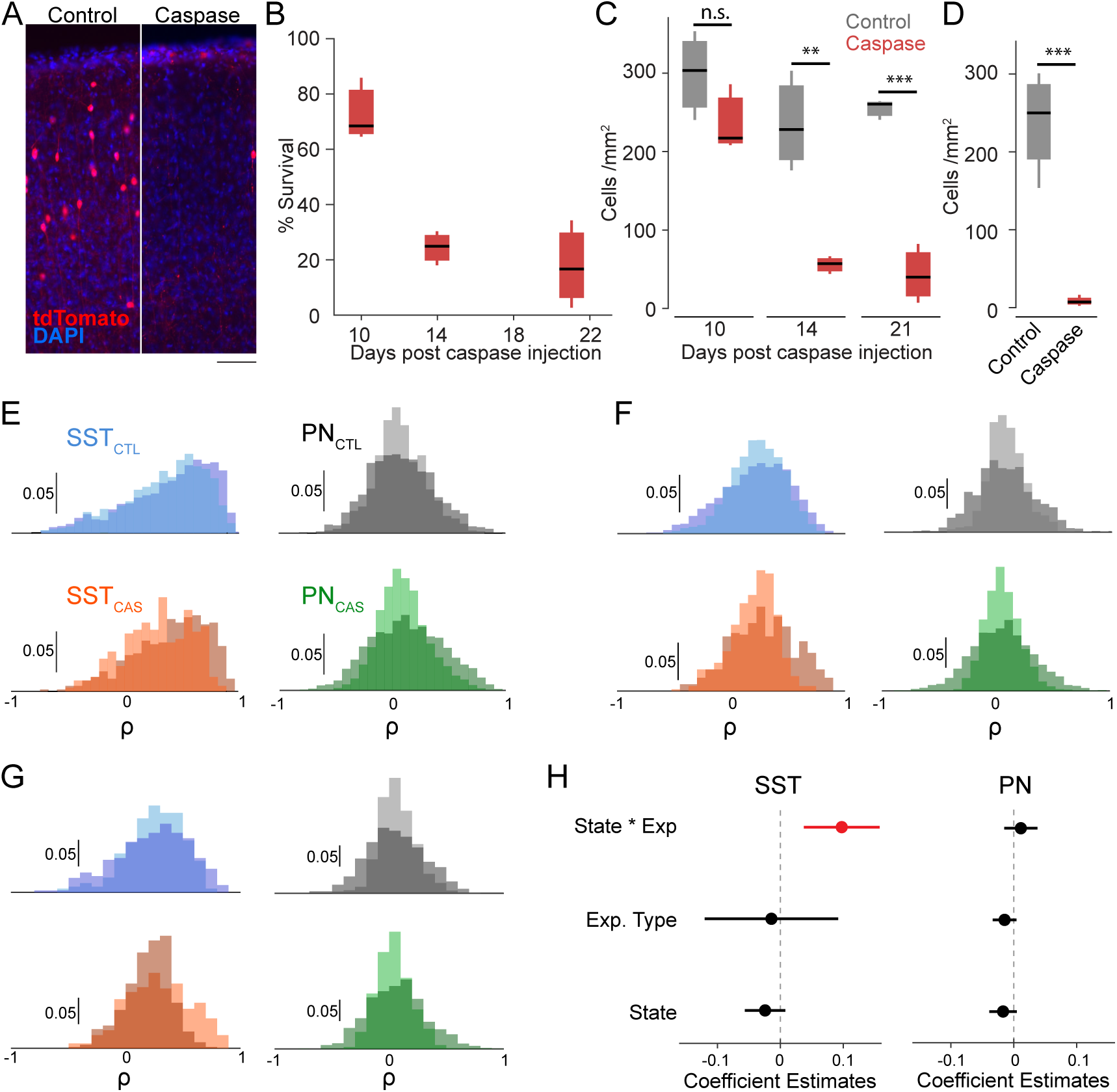
Ablation of VIP-INs by expression of caspase-3 and impact on the structure of local cortical activity. (A) Histology from an example VIP^Cre^Ai9^F/0^ mouse expressing tdTomato selectively in VIP-INs. VIP-INs (red) are present in V1 of the control hemisphere (left) but absent in the V1 injected with AAV-Syn-FLEX-taCasP3-TEVP. Scale bar = 70μm. (B) Percent survival of VIP-INs over time following caspase virus injection, calculated as [VIP-INs_caspase_/VIP-INs_control_] in each animal (n = 3 mice). (C) Density of VIP-INs in layer 2/3 of V1 cortex at 10, 14, and 21 days post caspase virus injection (n = 3 mice). (D) Density of VIP-INs in layer 2/3 of V1 cortex in animals used for 2-photon imaging (n = 4 control, 4 caspase mice). Unpaired t-test for histology, ** p<0.01, *** p<0.001. (E) Noise correlation distributions of ΔF/F_0_ for control (SST-INs, upper left, blue; PNs, upper right, grey) and VIP ablated (SST-INs, lower left, orange; PNs, lower right, green) animals. Correlations for sitting (lighter shades) and running (darker shades) are shown separately, with the overlap indicated by the darkest shade. SST control: n = 2180 pairs, 6 mice; SST VIP-ablated: n = 447 pairs, 4 mice; PN control: 1077 pairs, 5 mice; PN VIP-ablated: 2481 pairs, 6 mice. (F) Same as in E, but for deconvolved traces. SST control: n = 2180 pairs, 6 mice; SST VIP-ablated: n = 447 pairs, 4 mice; PN control: 1077 pairs, 5 mice; PN VIP-ablated: 2481 pairs, 6 mice. (G) Same as in E, but for mean-matched deconvolved data (see Methods). SST control: n = 1293 Q, 975 L pairs, 6 mice; SST VIP-ablated: n = 261 Q, 202 L pairs, 4 mice; PN control: 748 Q, 748 L pairs, 6 mice; PN VIP-ablated: 1658 Q, 1786 L pairs, 6 mice (H) Coefficient estimates of linear mixed effects model for noise correlations in G, with experiment type (control or VIP ablation), and state (quiescence or locomotion) with the interaction term (denoted state*exp) as fixed effects, and imaging field of view nested in the mouse as random effects. Horizontal bars for confidence intervals, red bar indicates significance (p<0.01).

**Supplementary Figure 2.**
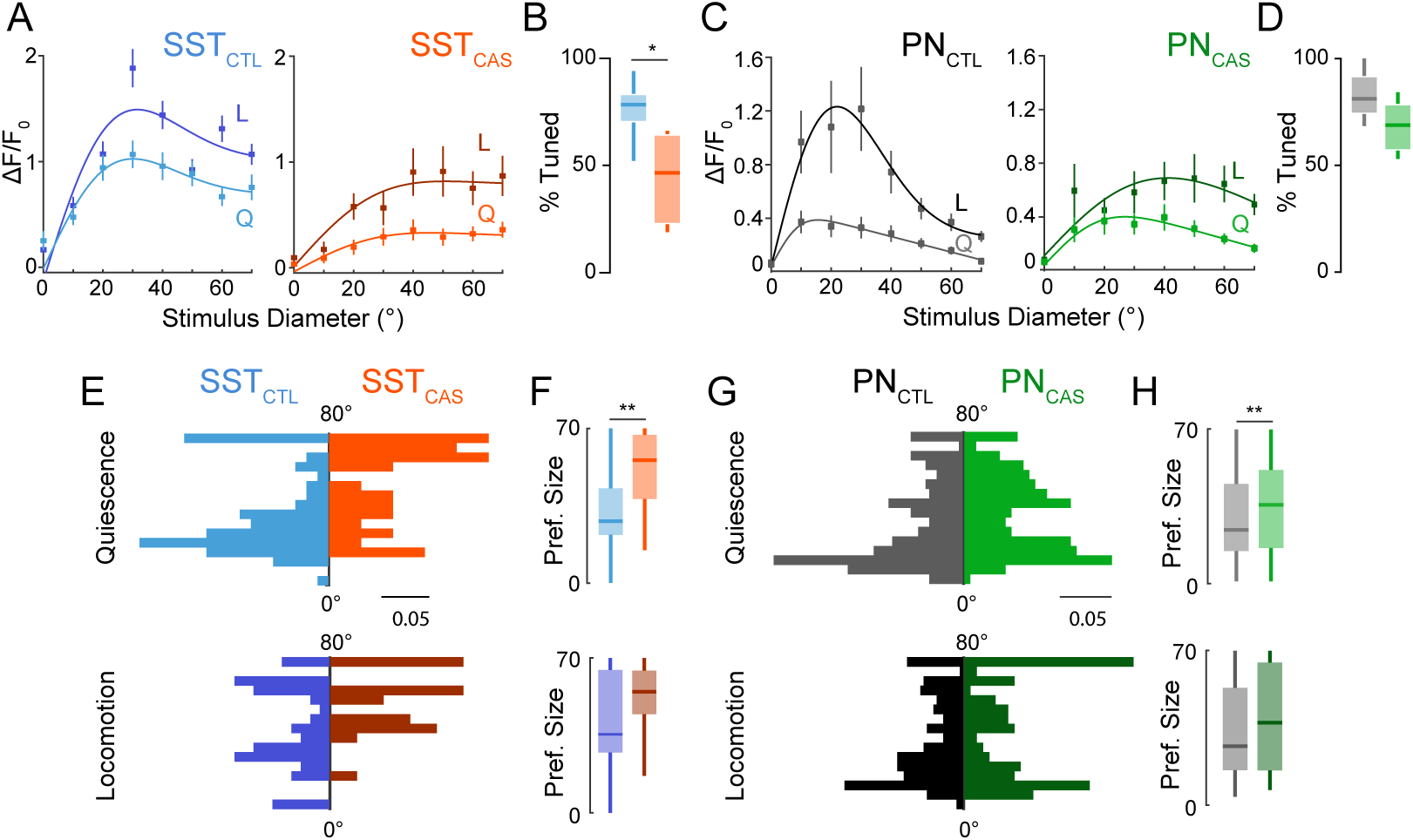
Preferred stimulus size for SST-INs and PNs. (A) ΔF/F_0_ visual responses of SST cells for periods of quiescence (Q, light lines) and locomotion (L, dark lines) for control (blue; SST_CTL_) and VIP ablation animals (orange; SST_CAS_). Baseline (F_0_) was set as the 1s period before the stimulus onset, where ΔF/F_0_= (F-F_0_)/ F_0_. (B) Boxplots of the percent of visually responsive cells that are visually tuned (see Methods) in SST control (blue; n= 6 mice) and VIP-ablation animals (orange; n= 4 mice). (C) Same as in A, but for PNs in control (gray; PN_CTL_) and VIP ablation (green; PN_CAS_) animals. (D) Same as in B but for PNs in control (gray; n = 6 mice) and VIP ablation (green; n= 5 mice) animals. (E) Probability distribution for the preferred stimulus size of SST-INs in control (blue; SST_CTL_) and VIP ablation (orange; SST_CAS_) animals. Distributions during quiescence are shown in light colors (upper; SST controls: n = 86 cells, 6 mice; SST VIP-ablation: n = 30 cells, 4 mice) and during locomotion in dark colors (lower; SST controls: n = 66 cells, 4 mice; SST VIP-ablation: 21 cells, 4 mice). (F) Boxplot of the population preferred stimulus size during quiescence (upper) and locomotion (lower). (G) Same as in E, for PNs in control (black; PNCTL) and VIP ablation (green; PNCAS) animals. (Quiescence, upper: PN controls: n = 254 cells, 6 mice; PN VIP-ablation: n = 175 cells, 5 mice. Locomotion, lower: PN controls: n = 202 cells, 3 mice; PN VIP-ablation: 122 cells, 3 mice). (H) Same as in F, for PNs. * p<0.05, ** p<0.01. Percent tuned: unpaired Student’s t-test. Preferred size: 0/1 inflated beta mixed effects regression model, with experiment type (control or VIP ablation) as fixed effect, mouse with nested imaging field of view as random effect.

**Supplementary Figure 3.**
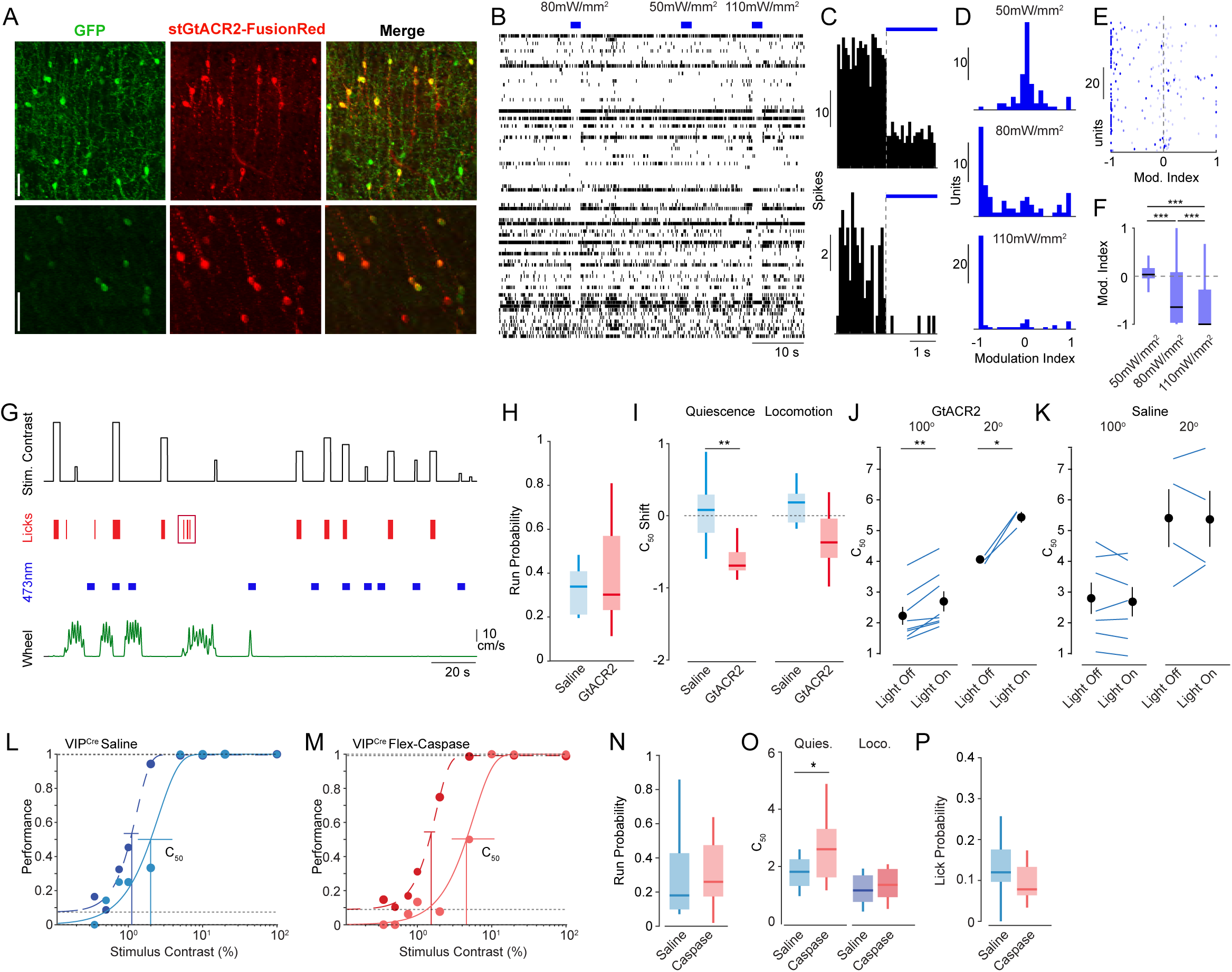
Modulation of perceptual behavior by manipulation of VIP-IN activity. (A) Example images of conditional GtACR2 expression (red) in GFP-expressing VIP-INs in V1 of an example animal (percent overlap: 79.8 ± 3.5; n = 3 mice). Scale bars = 50μm. (B) Raster plot of the spiking of 83 single neurons and multi-units in V1 cortex of a mouse expressing GtACR2 in excitatory pyramidal neurons. Light pulses (2 seconds duration) were given at 50, 80, or 110mW/mm^2^ to suppress firing. (C) Histograms of the firing of two example V1 neurons in response to pulses of blue light. (D) Histogram of the modulation index of recorded neurons in response to light pulses at 50, 80, or 110mW/mm^2^. (E) Raster plot of the modulation index for the 83 units, separated by light power. (F) Box plots of the population modulation index values of recorded neurons for 50, 80, or 110mW/mm^2^. *** p<0.001. Wilcoxon signed-rank test. (G) Schematic of events during one segment of an example contrast detection task session, including visual stimuli (black), lick responses (red), periods of bilateral illumination with 473nm light (blue), and wheel speed (green). False alarm lick responses are denoted by the red box. Height of the black trace denotes contrast from 0 to 100% on a log scale. (H) Box plots of probability of locomotion behavior in saline-treated (blue) and GtACR2-expressing (red) animals. (I) Change in perceptual threshold for contrast (C50) in response to activation of GtACR2 in VIP-INs during quiescence (left) and locomotion (right). (J) Raw C50 values for task periods with and without light activation of GtACR2 for 100° (left) and 20° (right) diameter stimulus sizes. Lines indicate values for individual animals. (K) Same as J, but for saline-treated animals. GtACR2: n = 8 mice for 100°, n = 3 mice for 20°. Saline: n = 7 mice for 100°, n = 4 mice for 20°. ** p<0.01, unpaired (I), paired (J,K) t-test. (L) Performance curves for an example saline-treated animal in the visual contrast detection task during quiescence (light blue) and locomotion (dark blue). (M) Same as in A, for an example VIP ablation animal. (N) Box plots of probability of locomotion behavior in saline-treated (blue) and VIP-ablated (red) animals. (O) C50 values for task performance during quiescence (left) and locomotion (right) in control (blue) and VIP ablation (red) animals. (P) False alarm rate for saline-treated and VIP ablation animals. VIP ablation: n = 8 mice. Saline: n = 14 mice. * p<0.05, Unpaired t-test.

## Notes

### Competing Interest Statement

The authors have declared no competing interest.

### Summary of Updates

Updated supplementary data plots and statistics.

